# Genome-wide CRISPR screens reveal multitiered mechanisms through which mTORC1 senses mitochondrial dysfunction

**DOI:** 10.1101/2020.10.22.351361

**Authors:** Kendall J. Condon, Jose M. Orozco, Charles H. Adelmann, Jessica B. Spinelli, Pim W. van der Helm, Justin M. Roberts, Tenzin Kunchok, David M. Sabatini

**Affiliations:** Whitehead Institute for Biomedical Research and Massachusetts Institute of Technology, Department of Biology, 455 Main Street, Cambridge, Massachusetts 02142, USA; Howard Hughes Medical Institute, Department of Biology, Massachusetts Institute of Technology, Cambridge, Massachusetts 02139, USA; Koch Institute for Integrative Cancer Research and Massachusetts Institute of Technology, Department of Biology, 77 Massachusetts Avenue, Cambridge, Massachusetts 02139, USA; Broad Institute of Harvard and Massachusetts Institute of Technology, 415 Main Street, Cambridge, Massachusetts 02142, USA

## Abstract

In mammalian cells, nutrients and growth factors signal through an array of upstream proteins to regulate the mTORC1 growth control pathway. Because the full complement of these proteins has not been systematically identified, we developed a FACS-based CRISPR-Cas9 genetic screening strategy to pinpoint genes that regulate mTORC1 activity. Along with almost all known positive components of the mTORC1 pathway, we identified many new genes that impact mTORC1 activity, including *DCAF7, CSNK2B, SRSF2, IRS4, CCDC43*, and *HSD17B10*. Using the genome-wide screening data, we generated a focused sublibrary containing single guide RNAs (sgRNAs) targeting hundreds of genes and carried out epistasis screens in cells lacking nutrient- and stress-responsive mTORC1 modulators, including GATOR1, AMPK, GCN2, and ATF4. From these data, we pinpointed mitochondrial function as a particularly important input into mTORC1 signaling. While it is well appreciated that mitochondria signal to mTORC1, the mechanisms are not completely clear. We find that the kinases AMPK and HRI signal, with varying kinetics, mitochondrial distress to mTORC1, and that HRI acts through the ATF4-dependent upregulation of both Sestrin2 and Redd1. Loss of both AMPK and HRI is sufficient to make mTORC1 largely resistant to mitochondrial dysfunction. Taken together, our data reveal a catalog of genes that impact the mTORC1 pathway and clarify the multifaceted ways in which mTORC1 senses mitochondrial dysfunction.

## Introduction

The mechanistic target of rapamycin complex 1 (mTORC1) is a eukaryotic cell growth regulator that responds to nutrient and growth factor availability. Under nutrient-replete conditions, mTORC1 licenses anabolic processes while inhibiting catabolic ones. Given the myriad of stimuli that mTORC1 responds to, it is no surprise that a diverse set of proteins, many as part of large complexes, act in a coordinated manner to regulate mTORC1 activity.

The heterodimeric Rag GTPases (RagA/B and RagC/D) play a central role in the control of mTORC1 by nutrients. In response to amino acids, as well as glucose and cholesterol, GTP-bound RagA/B and GDP-bound RagC/D mediate the recruitment of mTORC1 to the lysosomal surface (1–5). Once at the lysosome, GTP-bound Rheb, which is under the control of growth factors through the TSC complex pathway, binds to mTORC1 and stimulates its kinase activity (6–13). Together, Rheb and the Rags form a GTPase-based coincidence detector at the lysosomal surface that ensures that mTORC1 becomes activated only when nutrient and growth factor conditions are optimal. Given that the Rag and Rheb GTPases are central arbiters of mTORC1 activation the regulation of their respective nucleotide states is of great interest.

Dozens of proteins have been shown to modulate mTORC1 activity, many acting indirectly through one of several key effectors. However, the relative contributions of these proteins to the regulation of mTORC1 activity has not been systematically interrogated. Additionally, the majority of the proteins that regulate mTORC1 were identified using proteomic approaches. While fruitful, these studies leave open the possibility that proteins that play a role in mTORC1 regulation through transient or indirect interactions with pathway components, or are not easily detected with mass spectrometry-based proteomics, have not been identified.

Advances in CRISPR-Cas9-based screening have generated large catalogs of gene essentiality data in numerous cell lines, which can be leveraged to identify genes that are coessential with those encoding components of the mTORC1 pathway (14, 15). Though this type of analysis reveals many established mTORC1 regulators, a caveat is that it relies on cell fitness rather than mTORC1 activity as a read-out. A recent study utilized a gene-trap approach to identify mTORC1 regulators in haploid cells and define new relationships among established components (16). The CRISPR-screening strategy we present here expands this toolbox by enabling screening in a large set of genetically diverse cell lines of different lineages and allows for the identification of genes that regulate mTORC1 signaling but whose loss is not tolerated over the long term. We carried out a genome-wide CRISPR-Cas9 screen and a series of focused sublibrary screens to identify positive regulators of mTORC1. The hits from these screens ultimately led us to study how mTORC1 senses mitochondrial dysfunction. We find that two kinases (AMPK and HRI) act in a coordinated fashion to mediate the inhibition of mTORC1 caused by mitochondrial stress.

## Results

### A FACS-based genetic screen to identify mTORC1 regulators

To identify genes that contribute to mTORC1 activation, we devised a FACS-based CRISPR-Cas9 screening strategy that uses the phosphorylation of rpS6, a well-established marker of mTORC1 activity, as a readout (Fig.1*A*). We chose the HEK293T cell line for the screen because of its robust regulation of mTORC1 activity in response to amino acids and glucose starvation and restimulation (Fig.1*B*) and undertook initial screens in cells lacking both AMPKα1 and AMPKα2 (hereafter AMPK DKO cells) in order to more easily identify regulators that act independently of energetic stress (3). Generation of high-quality screening data from this strategy required careful consideration of downstream processing steps, so cell fixation, immunostaining, and DNA extraction protocols were extensively optimized.

**Fig. 1.**
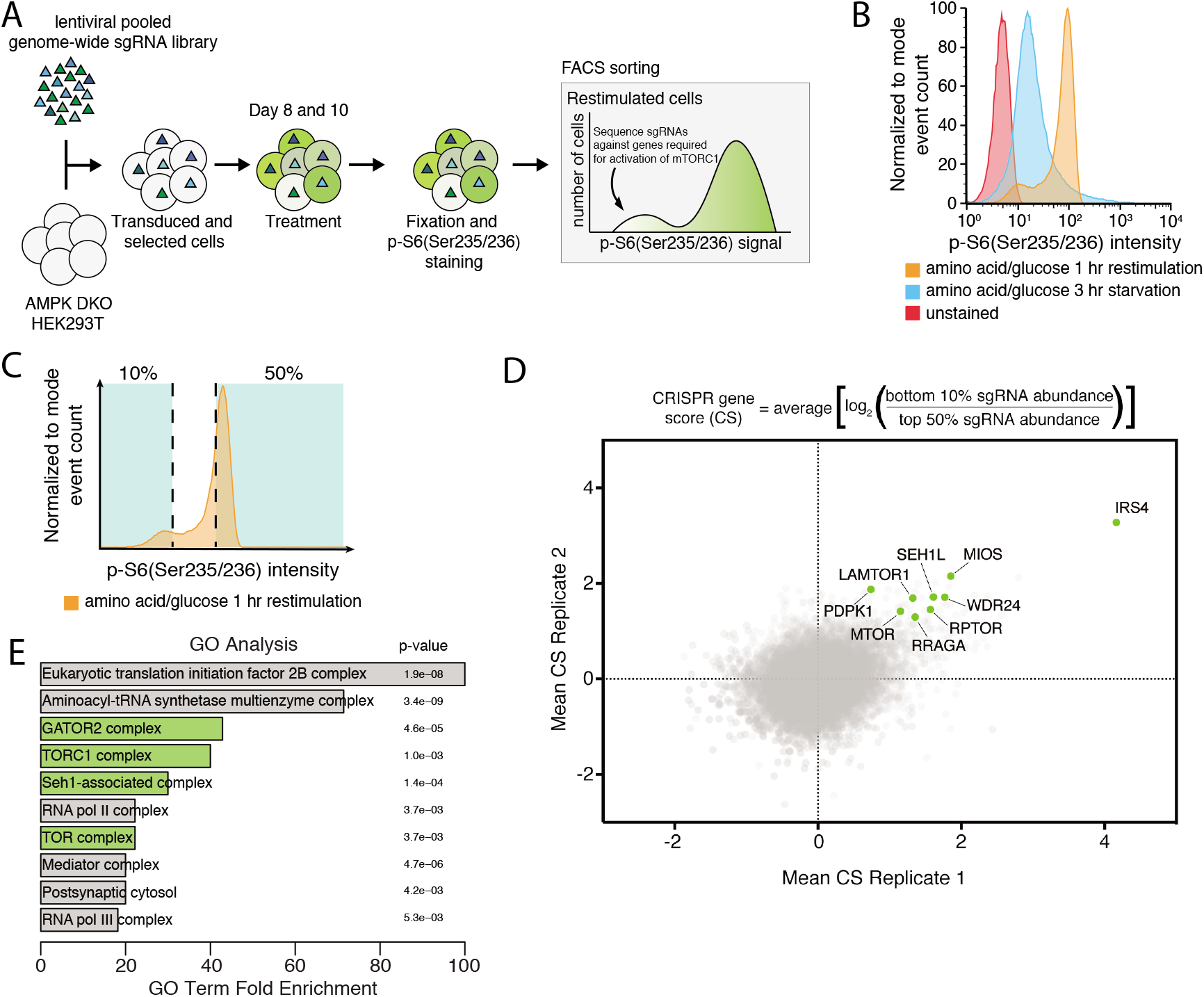
CRISPR-Cas9 screen identifies positive regulators of mTORC1. (A) Schematic of the CRISPR-Cas9 FACS-based genome-wide screen. (B) Representative flow cytometry histogram of wild-type HEK293T cells not exposed to the primary antibody (red), or immunostained after starvation of amino acids and glucose (blue) or starved of and restimulated with both (orange). (C) Schematic for FACS collection on a representative starved and restimulated sample of cells. (D) Equation for determining CRISPR scores (CS). Known mTORC1 regulators are highlighted in green. Positive CRISPR scores indicate genes that when lost prevented cells from fully reactivating mTORC1 upon combined glucose and amino acid restimulation as read-out by pS6 levels. Mean CS from two biological replicates of genome-wide screens in AMPK DKO HEK293T cells. Known positive regulators are highlighted in green. (E) GO analysis shows enrichment of mTORC1-related complexes in an unbiased manner. Analysis was performed on the 106 top scoring genes which had an FDR <.05.

AMPK DKO HEK293T cells were transduced with a lentiviral sgRNA library targeting ~18,000 genes and screened 8 and 10 days after to minimize the loss of sgRNAs targeting essential genes (Fig.1*A*). After immunostaining with a phospho-rpS6 antibody, cells were separated using a flow cytometry sorter into two fractions: those with the 10% lowest and 50% highest phopho-rpS6 signals (Fig. 1*C*). For each gene, a “CRISPR score” was generated by calculating the mean log2 fold-change in the abundances of sgRNAs targeting the gene between the two fractions. Replicate screens were compared directly to assess overlap and were also subjected to MAGeCK analysis (17). In this dataset, genes encoding central components of the mTORC1 pathway scored, including genes for the mTOR kinase (*MTOR*) and its associated protein raptor (*RPTOR*), along with established upstream regulators of mTORC1, including proteins in the nutrient sensing pathway such as the RagA GTPase (*RRAGA*) and Ragulator and GATOR2 subunits (*LAMTOR1, MIOS, SEH1L*, and *WDR24*) (Fig. 1*D*). Furthermore, gene ontology (GO) analysis confirmed in an unbiased fashion that the dataset is enriched for gene sets related to mTORC1 and amino acid signaling (Fig. 1*E*) (18). As expected because of their connection to the GCN2 pathway, which becomes activated when the levels of cytosolic uncharged tRNAs increase, genes encoding tRNA synthetases also scored highly in this analysis (19).

Overall, using a false discovery rate (FDR) cutoff of <0.1, our screening approach allowed us to identify 172 genes as potential regulators of mTORC1 activity, 76 of which have been previously defined as core fitness genes in the reference set from Hart et al. (20). This suggests that by performing the screens not long after library transduction, we were able to capture interactions between essential processes and mTORC1 signaling.

### Genes scoring in the genome-wide screen positively regulate mTORC1 signaling

For validation of hit genes from the screen, we used CRISPR-Cas9 to generate individual cell lines, each lacking a high scoring gene with a previously unknown or underappreciated link to mTORC1 signaling. Consistent with the screen, disruption of *DCAF7, CSNK2B, IRS4, CCDC43*, and *SRSF2/7* decreased phosphorylated rpS6 without impacting total rpS6 levels. To assess whether these genes act upstream of mTORC1, rather than impacting rpS6 phosphorylation through another mechanism, such as regulating an rpS6 phosphatase, we also examined the phosphorylation of S6 Kinase 1 (S6K1), which is a direct substrate of the mTORC1 kinase. Gratifyingly, when compared to the AAVS1 control sgRNA, loss of the hit genes decreased the phosphorylation of S6K1, suggesting that they likely act upstream of mTORC1 activity (Fig. 2*A*).

**Fig. 2.**
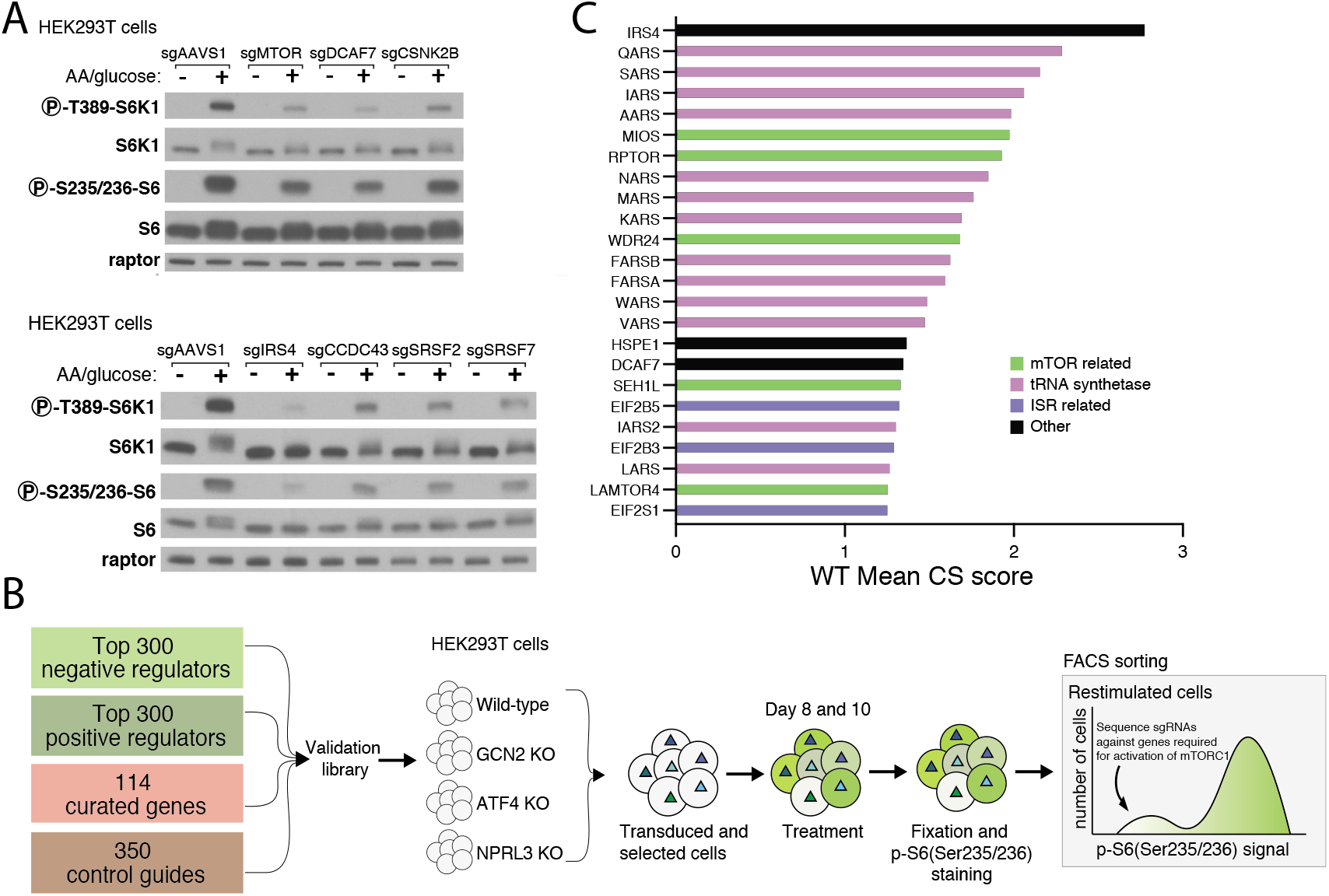
Generation of focused sublibrary and validation of individual gene hits. (A) Validation of select genes from screen. Immunoblot analysis shows mTORC1 signaling is blunted, as detected by decreased phosphorylation of the direct mTORC1 substrate S6K1, in HEK293T cells expressing sgRNAs targeting indicated genes. Signaling was assayed 8 days post-transduction with the indicated sgRNA as described for the primary screen. (B) Categories of genes included in the focused sublibrary along with a schematic of how the focused screens were performed. (C) CS scores for the 24 top scoring genes from a focused sublibrary screen in wildtype HEK293T cells.

In order to validate hits from the primary screen *en masse*, we generated a focused sublibrary of sgRNAs targeting ~700 genes identified from (1) the positive regulator screen described here, (2) prior screens aimed at discovering negative regulators of the mTORC1 pathway (unpublished), and (3) known regulators of the mTORC1 pathway that eluded detection in both screening strategies mentioned above (Fig. 2*B*). Screens with the focused sublibrary in wild-type HEK293T cells gave results that largely recapitulated those from the primary screens performed in the AMPK DKO cells, indicating that the screening strategy is robust and that the hits are not dependent on the loss of AMPK. Importantly, genes encoding known positive regulators of the mTORC1 pathway, including *MIOS, RPTOR, WDR24, SEH1L, LAMTOR2/4, RHEB, RRAGA*, and *MTOR*, all score within the top 24 genes (Fig. 2*C*).

### Focused sublibrary screens define epistatic relationships

In small-scale screens using the focused sublibrary, we asked whether hit genes impact mTORC1 through the integrated stress response (ISR) or the Rag-based amino nutrient-sensing pathway. To interrogate the role of the ISR, we used HEK293T cells lacking the GCN2 kinase, which is activated by uncharged tRNAs that accumulate upon amino acid starvation, or the ATF4 transcription factor, which can be induced by a number of kinases, including GCN2 (19, 21–24). To define genes acting through the Rag pathway, we generated HEK293T cells deficient in NPRL3, a component of GATOR1, a negative regulator of RagA/B whose loss renders mTORC1 signaling insensitive to amino acid or glucose starvation (25–27).

Comparisons of hits from sublibrary screens in wild-type HEK293T cells and those deficient in GCN2 or ATF4 showed that many of the tRNA synthetase genes did not score in the GCN2 or ATF4 KO cells (Fig. 3*A* and *B*), supporting a previous report that defects in tRNA charging impinge upon mTORC1 through GCN2 and ATF4 (28). Screens in the *NPRL3*-null HEK293T cells revealed two classes of genes that require GATOR1 to impact mTORC1 (Fig. 3*C*). The first consists of components of GATOR2, a known positive regulator of the pathway that likely acts upstream of GATOR1 to inhibit its function (25). The second consists of the aforementioned tRNA synthetases, supporting a previously proposed model in which defects in tRNA charging drive ATF4-dependent upregulation of Sestrin2, an inhibitor of GATOR2 (28–31).

**Fig. 3.**
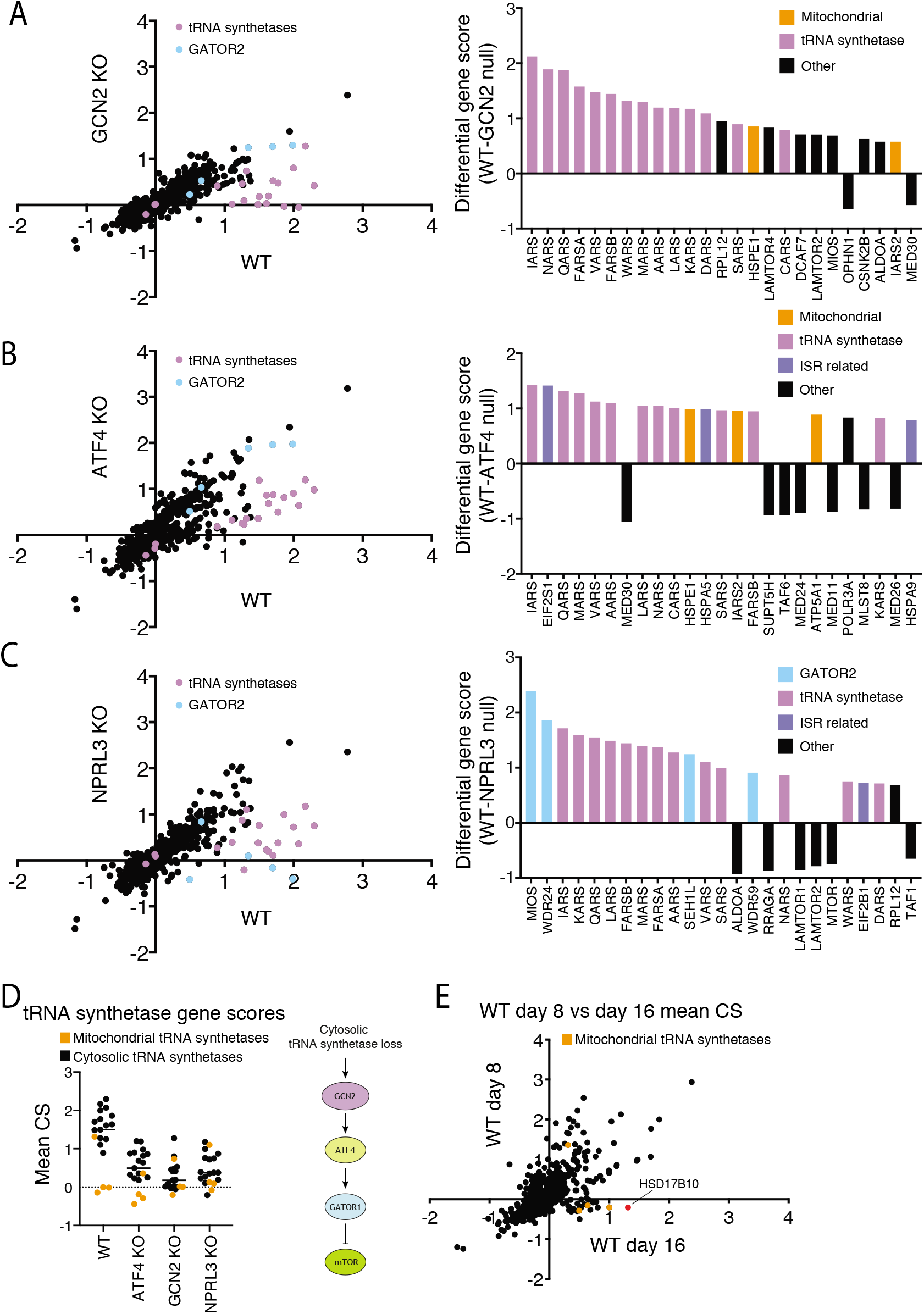
Focused sublibrary screens in cells deficient in known stress and amino acid sensing pathways. (A-C) Comparisons of CRISPR scores (CSs) from focused sublibrary screens in wild-type HEK293T to those in cells lacking GCN2, ATF4, or NPRL3. The top 25 differential CSs for each screening pair are included in an adjacent bar graph. The tRNA synthetase genes targeted in the library are highlighted in purple. Genes encoding GATOR2 components are highlighted in light blue. (D) Scores for tRNA synthetase genes present in the focused sublibrary in screens across all indicated cell lines. (E) Focused screens performed at 8 versus 16 days after transduction, highlighting genes whose scores are time-dependent.

Given that a variety of mitochondrial electron transport chain and ATP synthase inhibitors are known to dampen mTORC1 signaling (32–39), it was surprising that only the sgRNAs targeting the cytosolic, but not mitochondrial, tRNA synthetases scored as inhibiting mTORC1 (Fig. 3*D*). A potential explanation for this discrepancy is that it takes a significant amount of time for a defect in mitochondrial translation to reduce mitochondrially-encoded components of the electron transport chain to levels sufficient to cause mitochondrial dysfunction. Indeed, in focused sublibrary screens performed at longer time points after sgRNA transduction (16 days), the sgRNAs targeting the mitochondrial tRNA synthetases genes did score, as did sgRNAs targeting the *HSD17B10* gene, which encodes a key component of the mitochondrial RNase P complex required for tRNA processing (Fig. 3*E*) (40–42).

### Disruption of mitochondrial function inhibits mTORC1 signaling through AMPK and ATF4

Given that HSD17B10 was among the strongest hits from our focused screens, we asked how its loss might inhibit mTORC1. To validate it, we developed a conditional knockout system, in which doxycycline suppressed the expression of a cDNA encoding HSD17B10 in HEK293T cells lacking the endogenous *HSD17B10* gene (HSD17B10 dox-off cells). Gratifyingly, suppression of HSD17B10 strongly reduced mTORC1 signaling as read out by S6K1 phosphorylation (Fig. 4*A*). As expected, given its function, its loss also substantially decreased the levels of MT-ND1, which is encoded by the mitochondrial genome and thus depends on mitochondrial translation for its expression (Fig. 4*A*). In ATF4 KO but not AMPK DKO cells, mTORC1 was completely resistant to HSD17B10 loss (Fig. 4*A*), suggesting that chronic inhibition of mitochondrial protein synthesis suppresses mTORC1 signaling through ATF4 via the ISR.

**Fig. 4.**
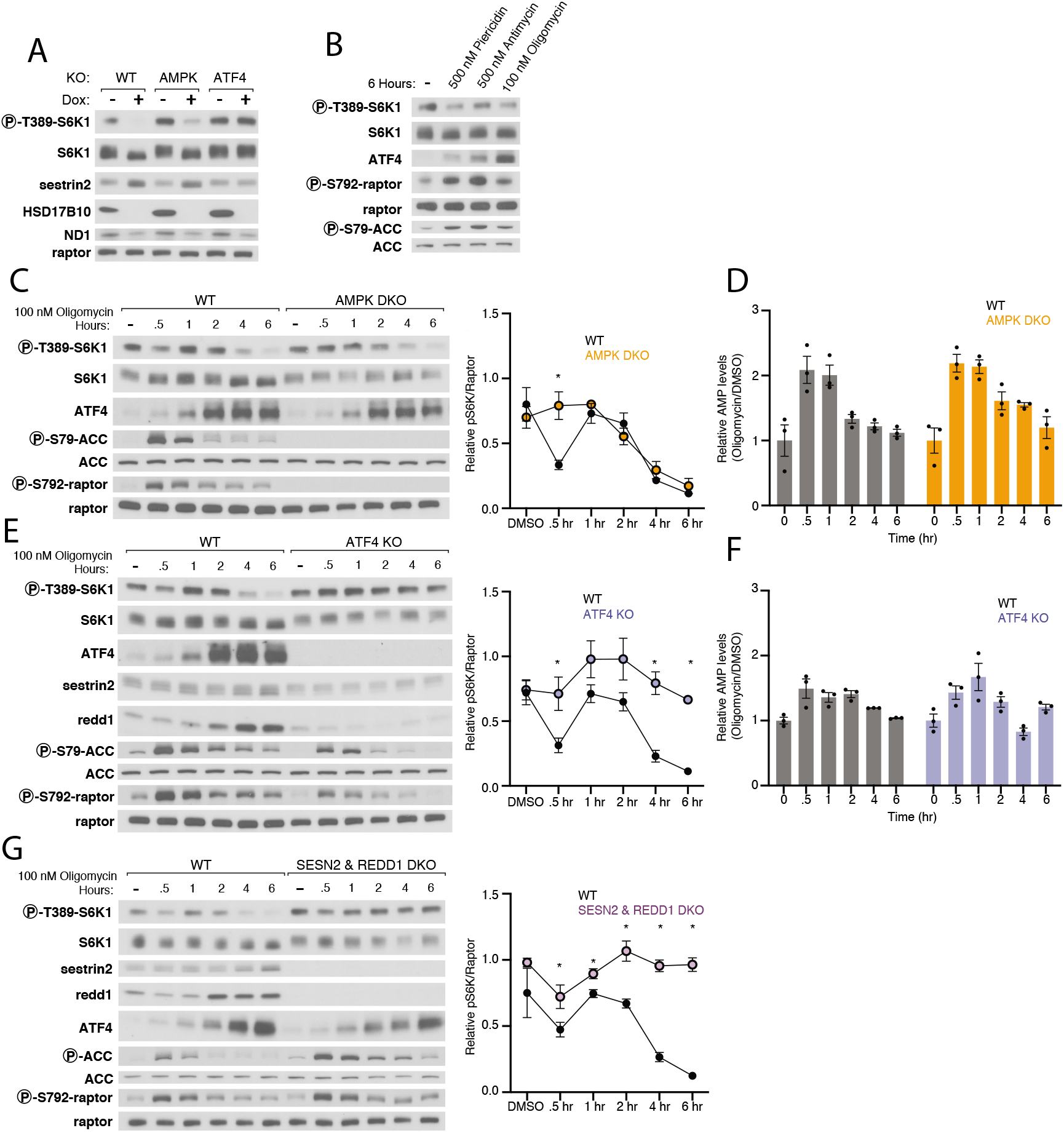
AMPK and ATF4 signal mitochondrial distress to mTORC1. (A) Loss of HSD17B10 inhibits mTORC1 activation by amino acids and glucose in an ATF4-dependent manner. HSD17B10 knock out HEK293T cells with dox-off conditional HSD17B10 expression were cultured with or without doxycycline (dox) for 8 days and then starved of amino acids and glucose for 1 hour and restimulated with both for 30 minutes. Immunoblot analyses of mTORC1 signaling and HSD17B10 expression. ND1 levels are expected to decrease upon loss of mitochondrial translation. (B) Inhibition of complex I, complex III, or the ATP-synthase suppresses mTORC1 signaling. mTORC1 activity was assayed in response to a 6-hour treatment with vehicle (DMSO), piericidin (500 nM), antimycin (500 nM), or oligomycin (100 nM). (C) AMPK mediates the first phase of mTORC1 inhibition in response to mitochondrial distress. Immunoblot analyses of mTORC1 signaling over the course of a 6-hour treatment with 100 nM oligomycin in wild-type and AMPK DKO HEK293T cells. The levels of S6K1 phosphorylation were quantified using densitometry with ImageJ. Values shown are the mean ± s.e.m. for *n*=3 biologically independent experiments. *P* values were determined using a two-sided Student’s *t*-test. \**P* < .05 (D) AMP levels increase acutely upon treatment with oligomycin and subsequently return to baseline levels. Metabolite extracts were analyzed by LC-MS and relative AMP levels are shown as the mean ± s.e.m. for *n*=3 biologically independent experiments. (E,F) ATF4 mediates the second phase of mTORC1 signaling in response to mitochondrial distress. Time course experiments were performed and analyzed as in (C,D). (G) Sestrin2 and Redd1 are required for the ISR to inhibit mTORC1 signaling in response to mitochondrial distress. Time course experiments were performed and analyzed as in (C,D).

Because mitochondrially-encoded proteins are components of the electron transport chain (ETC), we asked how its acute inhibition impacts mTORC1 signaling. We treated wild-type HEK293T cells with an inhibitor of complex I (piericidin), complex III (antimycin), or ATP synthase (oligomycin) for six hours. As expected, these treatments reduced the rate of mitochondrial oxygen consumption (*SI Appendix*, Fig. S1). The inhibitors also dampened mTORC1 signaling, activated AMPK, and induced ATF4 expression, with piericidin and antimycin having more mild effects than oligomycin (Fig. 4*B*). Given this, we carried out future experiments with oligomycin.

Over a 6-hour time course experiment in wild-type cells, oligomycin inhibited mTORC1 signaling in two distinct phases. A decrease in mTORC1 signaling first occurred after 30 minutes of oligomycin treatment, which correlated with an increase in AMP levels as well as AMPK activity, as detected by the phosphorylation of its substrates ACC and Raptor (Fig. 4*C* and *D*). After 1 hour of oligomycin treatment, mTORC1 signaling was restored to pre-treatment levels, but starting at 2-4 hours after oligomycin addition, mTORC1 was again inhibited but in this case in the absence of a substantial increase in AMP levels. Instead, at these later time points, the inhibition of mTORC1 correlated with an induction of the ISR as detected by an increase in ATF4 expression (Fig. 4*C*).

To understand the contributions of AMPK and ATF4 to the oligomycin-induced inhibition of mTORC1, we carried out oligomycin treatment time course experiments in the AMPK DKO and ATF4 KO cells. Although, at short time points AMP levels increased in both WT and AMPK DKO cells (Fig. 4*D*), in the absence of AMPK, mTORC1 signaling was not inhibited 30 minutes after addition of oligomycin (Fig. 4*C*). However, mTORC1 signaling in AMPK DKO cells remained sensitive to inhibition at longer time points (Fig. 4*C*), suggesting an AMPK-independent response was being activated.

In cells lacking ATF4, oligomycin did not inhibit mTORC1 signaling at either short or long time points after its addition (Fig. 4*E*). Indeed, in these cells oligomycin mildly increased mTORC1 signaling at the shorter time points, perhaps explaining why the AMPK-dependent inhibition of mTORC1 seen at 30 minutes after oligomycin addition in wild-type cells was absent in ATF4 KO cells. Importantly, loss of ATF4 did not greatly alter the impact of oligomycin on AMP levels or the phosphorylation of AMPK substrates (Fig. 4*E* and *F*). Thus, inhibition of the ATP synthase with oligomycin treatment inhibits mTORC1 in two phases, the first of which is dependent on AMPK and the second on ATF4.

### Both Sestrin2 and Redd1 are required for oligomycin to inhibit mTORC1 signaling

Given the critical role of ATF4 in mediating the inhibition of mTORC1 caused by oligomycin as well as loss of HSD17B10, we sought to understand the downstream mechanisms through which it acts. Mining of RNAseq datasets from cells with activated ATF4 revealed increases in the mRNAs for Sestrin2 and Redd1 (*DDIT4*), which encoded two well-known repressors of mTORC1 signaling (43–45). Sestrin2 is a leucine sensor that inhibits Rag-based nutrient signaling to mTORC1 by binding to GATOR2, while Redd1 inhibits growth factor signaling to mTORC1 via the Tuberous Sclerosis Complex (TSC) pathway (29, 30, 46, 47). It is well-appreciated that ISR induction via GCN2 activation upregulates Sestrin2, and Redd1 has been shown to be upregulated by PERK activation, which is also a component of the ISR (28, 48). In our hands, oligomycin increased Sestrin2 and Redd1 expression in a fashion that correlated with ATF4 levels and was dependent on ATF4 (Fig. 4*E*).

To test the roles of Sestrin2 and Redd1 in mediating the effects of oligomycin on mTORC1, we generated HEK293T cells lacking Sestrin2 or Redd1 or both. The individual loss of either Sestrin2 or Redd1 did not significantly impact the capacity of oligomycin to inhibit mTORC1 (*SI Appendix*, Fig. S2 *A* and *B*). In the Sestrin2 and Redd1 DKO cells (Fig. 4*G*), baseline mTORC1 signaling was slightly boosted, but it was still inhibited at 30 minutes of oligomycin treatment. At the later time points, however, loss of both Sestrin2 and Redd1, like that of ATF4, prevented oligomycin from inhibiting mTORC1. We conclude that induction of both Sestrin2 and Redd1 is required to inhibit mTORC1 signaling in response to oligomycin treatment at long time points. Moreover, there must be additional ATF4 targets that account for the boost in mTORC1 signaling seen in the ATF4 KO cells treated at early time points with oligomycin, as this effect was not prevented by the loss of Sestrin2 and Redd1.

### mTORC1 is largely insensitive to mitochondrial dysfunction in cells lacking the AMPK and HRI kinases

We next asked how oligomycin treatment induces ATF4 and inhibits mTORC1. ATF4 is induced at the level of its translation in a fashion that depends on the phosphorylation of the a-subunit of the translation initiation factor eIF2 (49, 50). Multiple kinases can phosphorylate eIF2α, including GCN2 in response to amino acid starvation and HRI in response to mitochondrial stress caused by oligomycin (19, 43, 45). In GCN2 KO cells, oligomycin still induced ATF4 and its targets Sestrin2 and Redd1, and also inhibited mTORC1(*SI Appendix*, Fig. S3). Thus, the relatively small impact that oligomycin has on amino acid levels like aspartate (*SI Appendix*, Fig. S4) is likely not sufficient to suppress tRNA charging and thereby activate GCN2.

In HRI KO HEK293T cells, mTORC1 remained sensitive to oligomycin at 30 minutes after its addition but was largely resistant to oligomycin at the longer time points (Fig. 5*A*). Consistent with this observation, HRI loss also prevented induction by oligomycin of ATF4 and its targets Sestrin2 and Redd1. Interestingly, in the HRI KO HEK293T cells, oligomycin increased AMPK signaling similarly to wild-type, but unlike in the wild-type cells, where it was quickly turned off, it remained active for the duration of the time course. This sustained AMPK signaling correlated with an inability of the HRI KO cells to restore normal levels of AMP following oligomycin treatment, indicating that maintenance of energy homeostasis upon inhibition of ATP synthesis requires not only AMPK but also HRI (Fig. 5*B*). Consistent with our finding that two kinases, AMPK and HRI, participate in the oligomycin-dependent inhibition of mTORC1, in HEK293T cells lacking both of them, mTORC1 was almost completely resistant to oligomycin treatment (Fig. 5*C*).

**Fig. 5.**
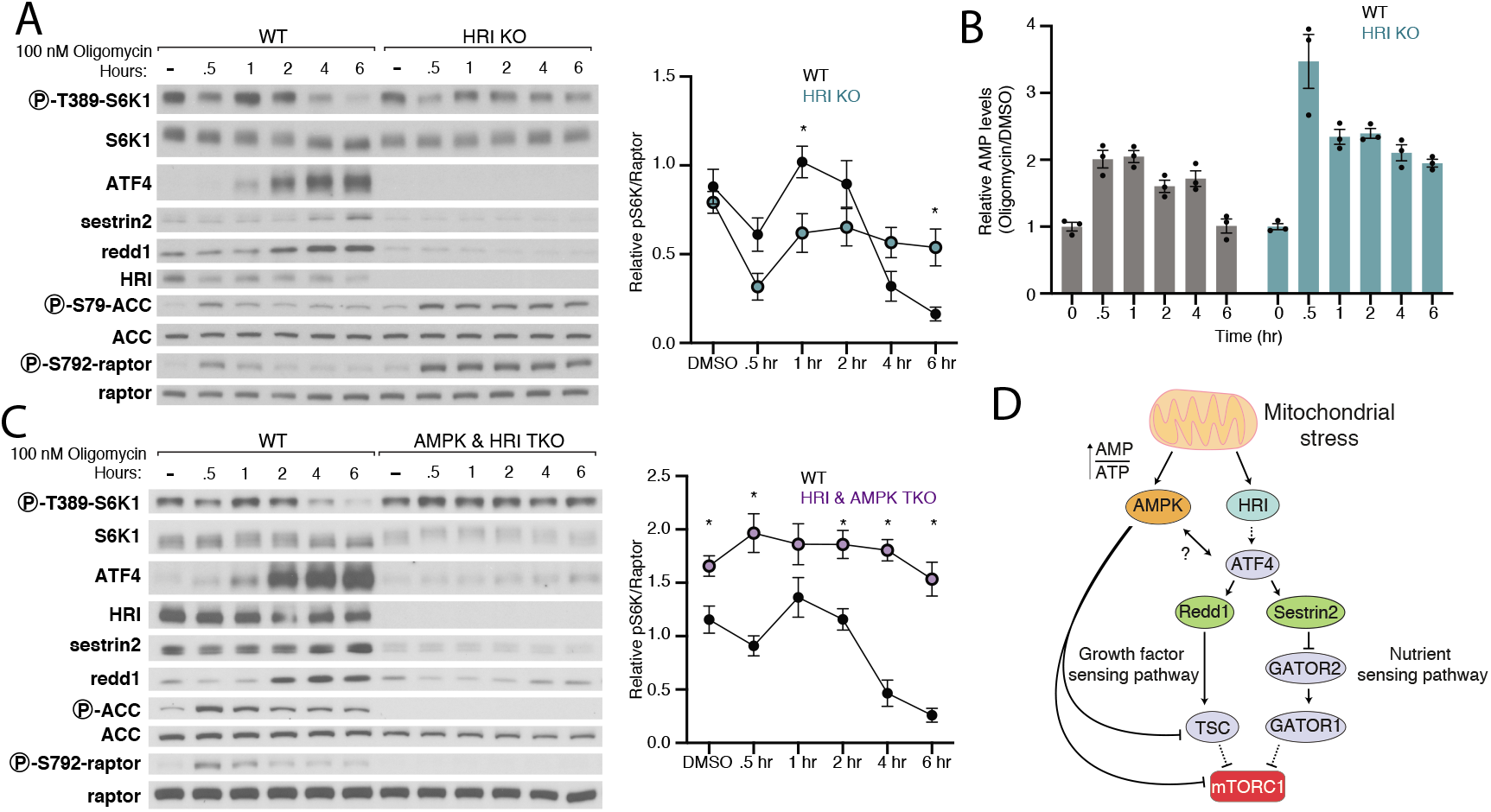
Together, AMPK and HRI signal mitochondrial dysfunction to mTORC1. (A) Loss of HRI prevents activation of the ISR in response to mitochondrial distress and renders mTORC1 signaling resistant to the second phase of inhibition caused by oligomycin. Cell lysates were analyzed by immunoblotting and the levels of S6K1 phosphorylation quantified as in Figure 4. Values are mean ± s.e.m. for *n*=3 biologically independent experiments. *P* values were determined using a two-sided Student’s *t*-test. (B) HRI KO HEK-293T cells have increased AMPK activity, which corresponds to an increase in AMP levels upon treatment with oligomycin. Metabolite extracts were analyzed by LC-MS and relative AMP levels are shown as the mean ± s.e.m. for *n*=3 biologically independent experiments. (C) Loss of both HRI and AMPK prevents inhibition of mTORC1 by oligomycin. Time course experiments were performed and analyzed as in (A,B). (D) Model for pathway leading to mTORC1 inhibition in response to mitochondrial distress.

## Discussion

Many upstream regulators control mTORC1 and facilitate its dynamic regulation by growth factors and nutrients. Given that dysregulation of the pathway is implicated in human diseases, such as cancer and neurodevelopmental disorders, there has been great interest in understanding how the pathway is regulated (51). The FACS-based CRISPR screens described here provide a systematic interrogation of the mTORC1 pathway in a human cell line in response to changes in nutrient levels. The genes identified in these screens will serve as a resource for future work and complement existing protein-protein interaction datasets.

In addition to identifying known positive regulators, we validated that the suppression of a number of genes not previously connected to mTORC1 can inhibit its activity, and these genes serve as leads for future investigation. For example, how does loss of *SRSF2 or SRSF7*, which encode splicing factors, impact mTORC1 signaling? A member of the same splicing factor family, SRSF1, has been previously implicated in mTORC1 signaling, but exactly how remains unclear (52). Other gene products that we identified, such as the kinase CSNK2B, might play roles in signaling cascades not previously connected to mTORC1 regulation. Our finding that loss of *IRS4* strongly inhibits mTORC1 activity is consistent with IRS4, rather than IRS1/2, being the main transducer of insulin signaling in HEK293-based cell lines, as previously reported (53). It is interesting to consider whether its tissue-specific expression might differentially modulate regulation of mTORC1 by insulin in vivo. Our focused sublibrary approach allowed us to efficiently perform many screens in different genetic backgrounds and to thus systematically define epistatic relationships between hundreds of potential hit genes and core stress pathways.

Finally, our study led us to concentrate on the relationship between mitochondrial stress and mTORC1. We find that two kinases, AMPK and HRI, but not GCN2, are necessary for signaling oligomycin-induced mitochondrial dysfunction to mTORC1 (Fig. 5*D*). AMPK inhibits mTORC1 at short time periods after mitochondrial inhibition while HRI does so at later times, so that loss of both kinases makes mTORC1 largely refractory to the effects of oligomycin. HRI acts through the transcriptional activation of two established mTORC1 pathway inhibitors, Sestrin2 and Redd1, via the ATF4 transcription factor. Indeed, loss of Sestrin2 and Redd1 together, but not of either alone, is sufficient to render mTORC1 resistant to the long-term inhibitory effects of oligomycin. Our findings also reveal that HRI is necessary for restoring normal levels of AMP upon prolonged oligomycin treatment and thus for inactivation of AMPK. This function of HRI is unlikely to require ATF4, as in cells lacking this transcription factor, AMP levels and AMPK activity responded largely normally to oligomycin treatment. Taken in sum, our work builds upon previous evidence that mitochondria “talk to” mTORC1 to bring new clarity to the mechanisms involved.

## Materials and Methods

### Cell culture

HEK293T cells were cultured in DMEM medium (Gibco) supplemented with 10% heat-inactivated FCS (Gibco) and penicillin-streptomycin. Cell lines were maintained at 37°C and 5% CO2. The mitochondrial response to stress is dependent on the cellular metabolomic state, so care should be taken to ensure cells are not overgrown or nutrient-starved ahead of experiments. All cell lines were obtained from ATCC (American Type Culture Collection) and tested for mycoplasma.

### Antibodies

The raptor antibody was from EMD Millipore (09-217); S6K1 pT389 (9234), S6K1 (2708), rpS6 pS235/S236 (2211), rpS6 (2217), Sestrin-2 (8487), ATF-4 (11815), raptor pS792 (2083), acetyl-CoA carboxylase (ACC) (3662), ACC pS79 (3661), and GCN2 (3302) antibodies were from Cell Signaling Technology (CST); ND1 (19703-1-AP), Sestrin2 (10795-1-AP), and Redd1 (10638-1-AP) antibodies were from Proteintech; HRI (MBS2538144) antibody was from MyBioSource; and the HSD17B10 (TA500724) antibody was from Life Technologies.

### sgRNA cloning, lentiviral production, and lentiviral transduction

Individual sgRNAs (Supplemental Table 1) were cloned into pLentiCRISPRv2 (Addgene #52961) at the BsmBI site as described by the depositor. Virus was generated as in (54). Briefly, 750,000 HEK293T cells were plated in DMEM supplemented with 20% IFS. At 12 hours post seeding, VSV-G envelope (Addgene #8454) and psPAX2 (Addgene #12260) were co-transfected with pLentiCRISPRv2 containing the indicated sgRNA sequences or pCW57 with the indicated cDNAs with XTremeGene 9 Transfection Reagent (Roche). After 12 hours, the culture medium was refreshed. At 28 hours post transfection, supernatant-containing virus was collected and passed through a 0.45 μm filter. Virus was stored at −80°C until use. For lentiviral transduction, cells were spinfected at 1,200g for 45 minutes at 37°C with 8 μg/mL polybrene and virus-containing medium. After 12 hours, the culture medium was refreshed, and after 24 hours, cells were selected with puromycin or blasticidin for 4 days.

### Pooled genome-wide CRISPR-Cas9 screens

The pooled genome-wide lentiviral sgRNA library (Addgene #1000000100) was prepared and screens were performed as described in (55) with slight modifications. Briefly, 600 million HEK293T cells were transduced with the viral pool to achieve 1000-fold library coverage after puromycin selection. After 72 hours of puromycin selection, 200 million cells were passaged and expanded every two days until day 7 when 1 billion cells were plated into fibronectin-coated 15 cm dishes. On days 8 and 10, the cells were washed three times with PBS supplemented with 1 mM CaCl2 and 500 μM MgCl2 and placed in RPMI lacking amino acids and glucose for 3 hours. After 3 hours, amino acids and glucose were added for 1 hour and the cells harvested. Cells were collected by washing once with ice-cold PBS, after which Accumax™ solution (Sigma) was added, and plates were nutated at 4°C for 10 minutes. After 10 minutes, cells were collected into a 50 mL centrifuge tube (Celltreat) and pelleted via centrifugation at 4°C at 300g for 3 minutes. Subsequently, cells were fixed by resuspension in formalin for 5 minutes at room temperature. At 5 minutes cells were pelleted via centrifugation at 4°C at 300g for 3 minutes. Finally, cells were permeabilized by resuspension in 90% methanol:PBS. After permeabilization cells were stored at −20°C until staining.

### Cell staining and fluorescence-activated cell sorting (FACS)

500 million previously fixed and permeabilized cells were pelleted by centrifugation at 4°C at 300g for 3 minutes and then washed once with ice-cold PBS. Cells were then blocked with 5% BSA in PBS for 1 hour at room temperature. For primary antibody staining, cells were incubated with primary antibody diluted (1:100) in 0.05% BSA in PBS at room temperature for 1 hour. After one wash with 0.05% BSA in PBS, cells were resuspended in secondary antibody diluted (1:200) in 0.05% BSA in PBS and incubated for 1hour. Finally, cells were washed twice in 0.05% BSA in PBS and filtered through a cell strainer prior to FACS sorting. Cells were sorted using a BD FACSAria III. Gates were drawn to collect the low intensity fraction at 10% and the high intensity fraction at 50%. Sorting was preformed until approximately 50 million cells were obtained for the lower fraction. Data were analyzed using FlowJo software (TreeStar).

### Analyses of CRISPR-Cas9 screens

Genomic DNA was extracted from sorted cells by first decrosslinking cells in a solution of 10 mM Tris pH 7.5 and 10 mg/mL Proteinase K (Roche) in PBS at 55°C for 24 hours. Subsequently, the Qiagen QIAmp DNA Blood Maxi kit was used according to the manufacturer’s instructions. High-throughput sequencing libraries were prepared as described in (56, 57). Sequencing reads were aligned to the sgRNA library and quantified. A pseudo count of 1 was added to each value and the log2-transformed fold change in abundance was then calculated between low and high fractions. MaGeCK analysis was performed as in (58) and (17).

### Focused sgRNA sublibrary screens

The focused sublibrary (Supplemental table 2) was designed by selecting the top scoring genes in the positive regulator screen described here and prior screens aimed at discovering negative regulators of the mTORC1 pathway (unpublished). Additionally, a curated list of known mTORC1 related genes and control sgRNAs were included. 10 sgRNAs per gene were selected from (Addgene #1000000100) along with 350 control sgRNAs that were selected randomly from this library. The oligonucleotide pool was synthesized by Agilent Technologies and cloned as described in (57). Focused sublibrary screens were carried out as described for genome-wide screens with modifications. Briefly, 60 million wild-type or indicated knockout HEK293T cells were transduced with the viral pool to achieve 1000-fold library coverage after puromycin selection. After 72 hours of puromycin selection, 20 million cells were passaged and expanded every two days until day 7, when 100 million cells were plated into fibronectin-coated 15 cm dishes. Screen replicates were analyzed by averaging high and low ratios and then calculating mean CS.

### Gene Ontology (GO) Term Enrichment

GO term enrichment was performed in R using the topGO R-package (18). Top scoring positive regulator genes with an FDR less than 5% from MAGeCK analysis were compared to all genes in the genome-wide library using the classic Fisher statistic from the topGO package for GO terms in the cellular component category. GO-terms with less than 3 members were filtered out by setting the node size to 3. Terms with p-values greater than 0.01 were filtered out and then ranked by fold enrichment.

### Starvation, drug treatments, and time course experiments

Cells were seeded at 720,000 cells per well in a fibronectin-coated six-well plate the day before an experiment. For nutrient starvation experiments, cells were washed once with RPMI lacking amino acids and glucose and then starved in 2 mL of the same media. After 1-hour, amino acids and glucose were added back for 30 minutes at which point cells were harvested and lysed. For drug treatments and time course experiments, RPMI containing DMSO, oligomycin (100 nM), antimycin (500 nM), or piericidin (500 nM) was prepared and medium was refreshed at indicated time points. Cells were collected simultaneously.

### Cell lysis

Cells were briefly washed with ice-cold PBS and then scraped into a Triton-X-100-based lysis buffer (1% Triton X-100, 10 mM beta-glycerol phosphate, 10 mM pyrophosphate, 40 mM HEPES pH 7.4, 2.5 mM MgCl2 with 1 tablet of EDTA-free protease inhibitor (Roche) per 50 mL buffer). Cell lysates were clarified by centrifugation at 17,000g at 4°C for 10 minutes.

### Generation of the dox-off and knockout (KO) HEK293T-based cell lines

The cDNA for human *HSD17B10* was codon optimized and synthesized by IDT as a gBlock™ Gene Fragment. This gene fragment was then cloned into the pCW57.kc1 (Addgene #pending) vector using NEBuilder^®^ HiFi DNA Assembly (NEB). Cells were transduced with lentivirus generated with the pCW57.kc1 vector and selected with blasticidin for 3 days. Subsequently, KO cells were generated using either px330 (Addgene #42230) or pLentiCRISPRv2 (Addgene #52961). After transfection as described in (3) or transduction, cells were selected with puromycin for 4 days and then allowed to recover for 7 days. Cells were then sorted by a flow cytometer into 96-well plates containing 200 μL DMEM medium (Gibco) supplemented with 30% heat-inactivated FCS (Gibco) and penicillin-streptomycin in each well. Clones were screened for KO by immunoblotting for proteins encoded by targeted genes. Double and triple KO cell lines were generated with the same process, beginning with SESN2 KO or AMPK DKO. AMPK DKO, SESN2 KO, and ATF4 KO HEK293T cells were generated as described in (3), (30), and (44), respectively.

### Treatment with doxycycline

Doxycycline was prepared as a stock solution (30 μg/mL), and stored at −80°C. Cells were cultured in DMEM medium (Gibco) supplemented with 10% heat-inactivated FCS (Gibco) and penicillin-streptomycin and 30 ng/mL Doxycycline for 10 days prior to experiments.

### Western blot quantifications

Western blot band intensities were measured using ImageJ. Average pixel intensities were collected over equal area boxes for bands and nearby areas without bands as background controls. Pixel densities were then inverted by subtracting 255 from all values. Background controls were then subtracted from corresponding band intensities. Finally, net pS6K values were divided by net raptor values. Because the mobility of S6K changes depending on its phosphorylation state, which can confound analysis, we chose to normalize pS6K levels to the raptor loading control.

### Oxygen consumption rates

Oxygen consumption rates (OCR) were measured using a Seahorse XFe96 Analyzer (Agilent). 25,000 cells were seeded 18 hours prior to the experiment in fibronectin-coated Seahorse XFe96 cell culture plates. 1 hour prior to the experiment, cells were incubated in Seahorse Media (Agilent) supplemented with 10 mM glucose, 2 mM glutamine, and 1 mM pyruvate containing vehicle (DMSO), oligomycin (1 μM), antimycin (500 nM), or piericidin (500nM). Three OCR measurements were taken for each treatment.

### Preparation of samples for metabolomics

Cells were washed twice with ice cold PBS and extracted on dry-ice in 0.8 mL 80% methanol containing 500 nM internal standards (Metabolomics Amino Acid Mix Standard: Cambridge Isotope Laboratories, Inc.). Cell extracts were collected using a cell scraper and transferred to a microcentrifuge tube. Samples were vortexed for 10 minutes at 4°C and centrifuged at 17,000g for 10 minutes at 4°C. Supernatants were transferred to new microcentrifuge tubes and evaporated to dryness by vacuum centrifugation. Dried polar extracts were stored at −80°C until analysis.

### Metabolomics

Metabolite profiling was conducted on a QExactive bench top orbitrap mass spectrometer equipped with an Ion Max source and a HESI II probe, which was coupled to a Dionex UltiMate 3000 HPLC system (Thermo Fisher Scientific, San Jose, CA). External mass calibration was performed using the standard calibration mixture every 7 days. Typically, dried polar fractions were reconstituted in 100 uL water and 2 uL were injected onto a SeQuant^®^ ZIC^®^-pHILIC 5μm 150 x 2.1 mm analytical column equipped with a 2.1 x 20 mm guard column (MilliporeSigma). Buffer A was 20 mM ammonium carbonate, 0.1% ammonium hydroxide; Buffer B was acetonitrile. The column oven and autosampler tray were held at 25°C and 4°C, respectively. The chromatographic gradient was run at a flow rate of 0.150 mL/min as follows: 0-20 min: linear gradient from 80-20% B; 20-20.5 min: linear gradient from 20-80% B; 20.5-28 min: hold at 80% B. The mass spectrometer was operated in full-scan, polarity-switching mode, with the spray voltage set to 3.0 kV, the heated capillary held at 275°C, and the HESI probe held at 350°C. The sheath gas flow was set to 40 units, the auxiliary gas flow was set to 15 units, and the sweep gas flow was set to 1 unit. MS data acquisition was performed in a range of *m/z* = 70–1000, with the resolution set at 70,000, the AGC target at 1×10^6^, and the maximum injection time at 20 msec. An additional scan (*m/z* = 220-700) was included in negative mode only to enhance detection of nucleotides. Relative quantitation of polar metabolites was performed with TraceFinder™ 4.1 (Thermo Fisher Scientific) using a 5 ppm mass tolerance and referencing an in-house library of chemical standards.

## Acknowledgments

We thank all members of the Sabatini lab for helpful insights; G. Liu and P. Rosen for critical reading of the manuscript; C. Thoreen (Yale) for the gift of the ATF4-null HEK293T cell line; and P. Thiru for help with MAGeCK analysis. We thank the Metabolite Profiling Core Facility at the Whitehead Institute for running metabolomics samples and for data analysis. This work was supported by grants to D.M.S. from the National Institutes of Health (NIH; R01 CA103866, R01 CA129105 and R01 AI047389). K.J.C. was supported by an MIT School of Science Fellowship in Cancer Research and an NSF fellowship (2016197106). J.M.O. was supported by a fellowship (F30CA210373) from the National Cancer Institute and the Harvard–MIT MSTP training grant (T32GM007753) from the National Institute of General Medical Sciences. C.H.A had fellowship support from the NIH (NRSA F31 CA228241-01). J.B.S was funded by the National Cancer Institute F99/K00 predoctoral to postdoctoral transition fellowship (K00CA234839). J.M.R. was supported by a fellowship grant from the National Cancer Institute of the National Institutes of Health (F31CA232355). D.M.S. is an investigator of the Howard Hughes Medical Institute and an American Cancer Society research professor.

## Author Contributions

K.J.C. and D.M.S. designed the research plan; K.J.C. performed the majority of the experiments with the aid of J.M.O., J.B.S., P.W.H., and T.K.; K.J.C., J.M.O. and C.H.A. contributed new reagents/analytical tools; K.J.C., J.M.O. and J.M.R. analyzed data, and K.J.C. wrote and K.J.C. and D.M.S. edited the paper.

---

**Fig. S1.**
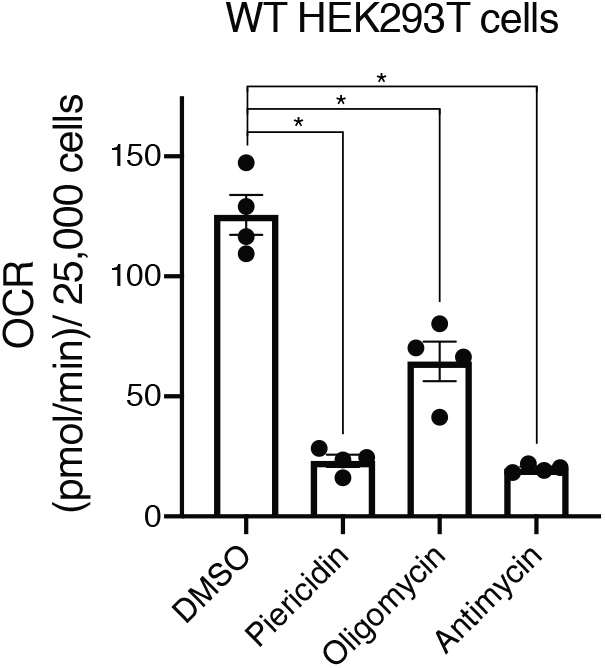
Mitochondrial inhibition with piericidin, antimycin, or oligomycin, lowers oxygen consumption rate (OCR). Wild-type HEK293T cells were treated with vehicle (DMSO), piericidin (500 nM), oligomycin (1 μM), or antimycin (500 nM) and OCRs were measured with a Seahorse XFe96 Analyzer. OCRs are shown as mean ± s.e.m. for *n*=4 biologically independent experiments. *P* values were determined using a two-sided Student’s *t*-test. \**P* < .05.

**Fig. S2.**
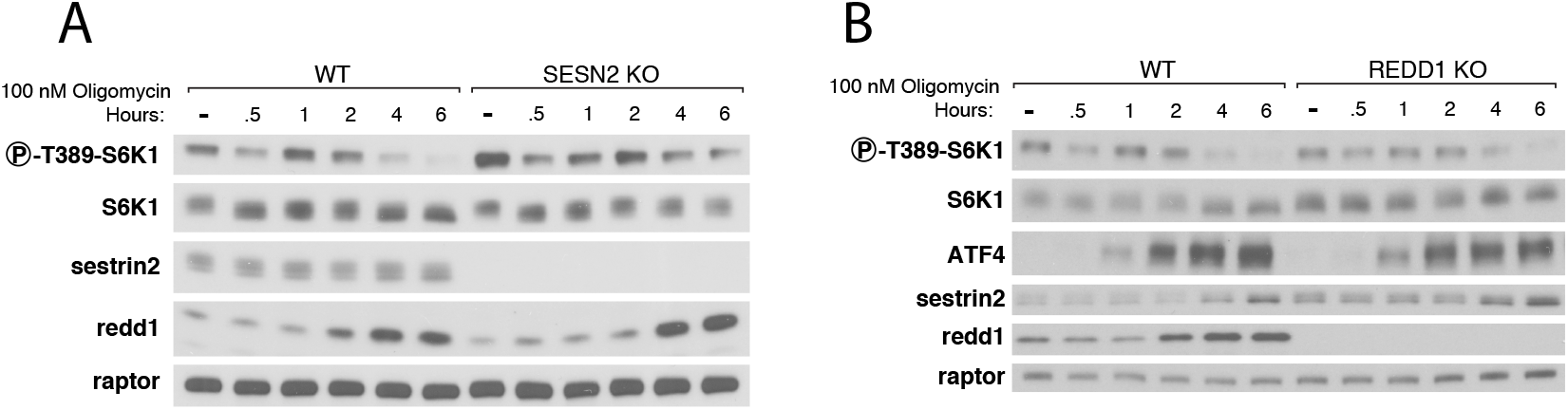
Loss of Sestrin2 or Redd1 alone is not sufficient to render mTORC1 signaling resistant to oligomycin treatment. (A) Immunoblot analyses of mTORC1 signaling over the course of a 6-hour treatment with 100 nM oligomycin in wild-type and Sestrin2 KO HEK293T cells. (B) Immunoblot analyses of mTORC1 signaling over the course of a 6-hour treatment with 100 nM oligomycin in wild-type and Redd1 KO HEK293T cells.

**Fig. S3.**
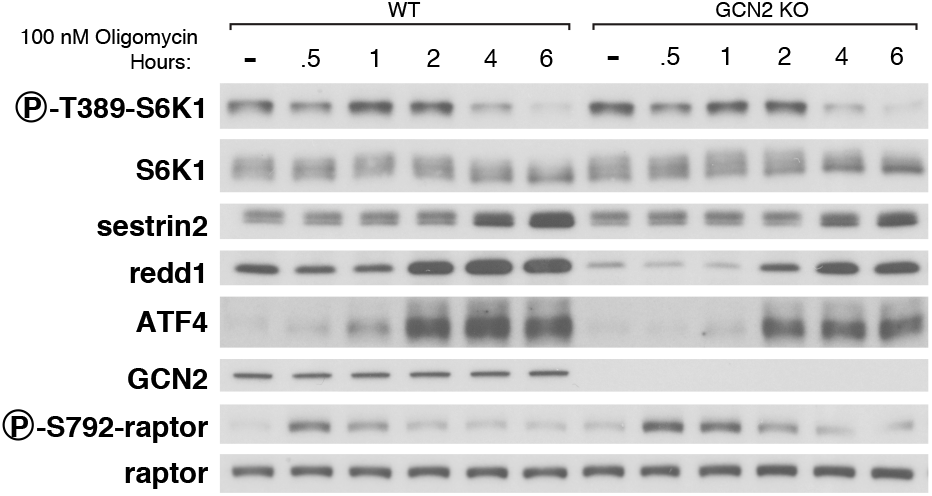
mTORC1 signaling remains sensitive to oligomycin in cells lacking GCN2. Immunoblot analyses of mTORC1 signaling over the course of a 6-hour treatment with 100 nM oligomycin in wild-type and GCN2 KO HEK293T cells.

**Fig. S4.**
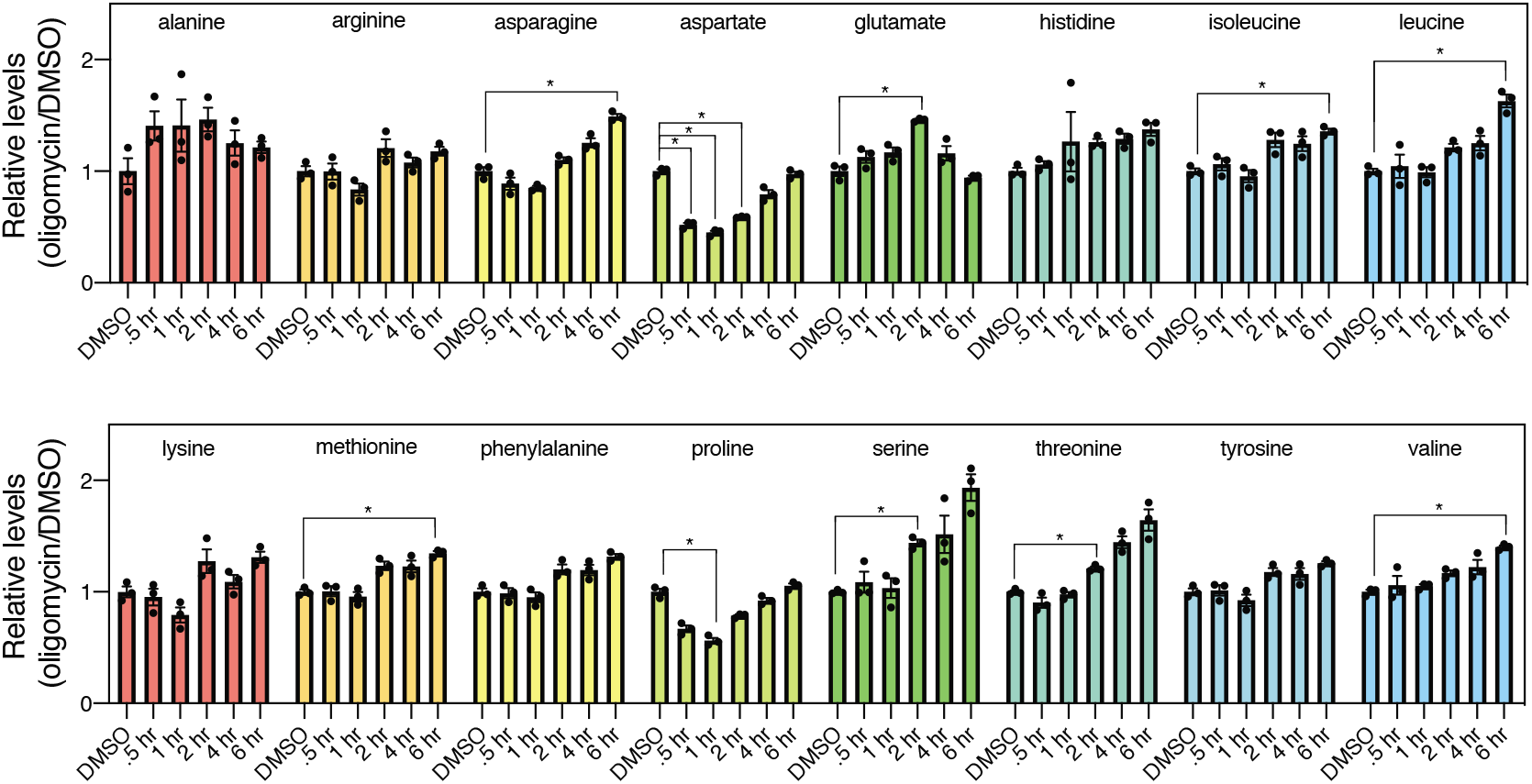
The levels of most amino acid do not decrease after a 6-hour treatment with oligomycin. Relative amino acid levels in wild-type HEK293T cells after a 6-hour treatment with 100 nM oligomycin. Relative amino acid levels are shown as mean ± s.e.m. for *n*=3 biologically independent experiments. \**P* <0.05 (\**P* <0.000625 after Bonferroni correction).

